# Autoencoder networks extract latent variables and encode these variables in their connectomes

**DOI:** 10.1101/2020.03.04.977702

**Authors:** Matthew Farrell, Stefano Recanatesi, R. Clay Reid, Stefan Mihalas, Eric Shea-Brown

## Abstract

Spectacular advances in imaging and data processing techniques are revealing a wealth of information about brain connectomes. This raises an exciting scientific opportunity: to infer the underlying circuit function from the structure of its connectivity. A potential roadblock, however, is that – even with well constrained neural dynamics – there are in principle many different connectomes that could support a given computation. Here, we define a tractable setting in which the problem of inferring circuit function from circuit connectivity can be analyzed in detail: the function of input compression and reconstruction, in an autoencoder network with a single hidden layer. Here, in general there is substantial ambiguity in the weights that can produce the same circuit function, because largely arbitrary changes to “input” weights can be undone by applying the inverse modifications to the “output” weights. However, we use mathematical arguments and simulations to show that adding simple, biologically motivated regularization of connectivity resolves this ambiguity in an interesting way: weights are constrained such that the latent variable structure underlying the inputs can be extracted from the weights by using nonlinear dimensionality reduction methods.

## 1 Introduction

The past years have seen spectacular effort, and spectacular success, in mapping synaptic connections in the brain. For example, the hemi-brain connectome of *Drosophila* was recently released, and imaging of the whole brain connectome is currently underway [1]. The synaptic connections in a cubic millimeter of mouse visual cortex have also recently been imaged [2]. These data build on pioneering efforts to map the connectome of *c. elegans*. From these advances there have emerged stunning new opportunities and new challenges for modelers and theoreticians.

Connectivity data has long been leveraged to shed light on circuit function. This includes the discovery of hierarchical organization in mammalian visual systems [3], which is thought to support the assembly of complex and abstract visual information in higher areas out of combinations of simple, local neural responses in lower areas; and the repeated structure across different neocortical regions [4, 5], indicating that neocortex supports general-purpose learning. As connectivity data become more complete, they can be used to more precisely constrain models [6]. For instance, while a variety of mechanisms have been proposed to explain motion processing in the retina, recent connectivity data allowed the authors of [7] to refine this class of models into one that better fits the observed connectivity. Additional studies link connectivity to the function of sensory circuits through the lens of increasing or decreasing the dimension of their inputs. For example, connections between mossy-fiber and granule cells in the *Drosophila* mushroom body appear to be random and sparse [8], which is thought to support associative learning by expanding dimension [9–11]. Other studies point out physical bottlenecks in sensory circuits that strongly suggest a compression of dimension, particularly in early visual pathways (for a review, see [12]). Such a compression forces circuits to select elements of their inputs which are necessary for downstream computations. Often this operation is modeled as extracting a low-dimensional set of *latent variables* that generate the higher-dimensional input signal.

Our work follows on these observations by probing the following question: in compressive circuits, can the actual structure of the selected-for latent variables be extracted from the connectome? While this question will prove to be a significant challenge to answer in general, here we start with a simple and mathematically tractable model of input compression: a linear autoencoder with a single layer of hidden units. This work sets out to discover if the weights that optimally compress inputs (in an L2 reconstruction sense) contain recoverable information about the input stimulus; namely, the latent variables generating the inputs. Here we focus on using dimensionality reduction methods on the weight matrices to recover this information.

Our main finding is that structure from the latent variables underlying the inputs *can* be extracted from the weights of our network, provided that the model is regularized by biologically inspired costs on weight resources (i.e. penalizing large weights). The structure that can be extracted includes the basic topology of the latent space; in particular, it can be inferred if the latent variables live on a space that wraps around (like a circle) or that doesn’t wrap around (like a line). We give both mathematical arguments in the case of linear autoencoders and the simulation results of training autoencoders with hyperbolic tangent nonlinearities. Our results have several important implications. First, they are an important proof of concept that meaningful information about network function can be extracted from the optimal weights alone. Second, they shed light on *how* the weights can be processed to reveal this latent structure. Specifically, we show that nonlinear (as opposed to linear) dimensionality reduction techniques are important for finding this structure. This can guide efforts to find structure in the weights of more complex models or biological data. Third, we establish that combinations of latent variables are reflected in the weight structure in a predictable way. This can be used as a tool for looking at complicated inputs that may be formed from many latent variables. These results reinforce an emerging narrative in the analysis of neural networks: that regularization is important not only for producing network models that generalize, but also for producing models that are interpretable (see [13, 14]).

## 2 Network Architecture

The model we consider is an autoencoder network with a single layer of hidden units, as illustrated in Fig. 1. The equation for the hidden unit activations in response to an input ***x***_*s*_ is

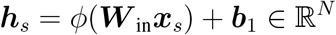

where *N* is the number of hidden units and *ϕ* is the activation function for the hidden units. The length *N* vector ***b***_1_ is a bias term. For our mathematical analysis we take *ϕ* equal to the identity, but we also show the results of simulations where *ϕ* = tanh. The activation of the output units is

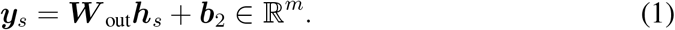

where *m* is the dimension of the input space. The network is trained via stochastic gradient descent (SGD) with momentum (RMSprop) to minimize the regularized L2 reconstruction error

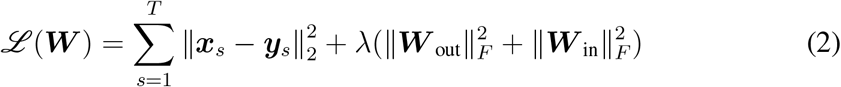

where *T* is the size of the training dataset. Here the network is trained so that outputs ***y***_*s*_ are close to ***x***_*s*_, regularized by a cost on weights.

**Figure 1:**
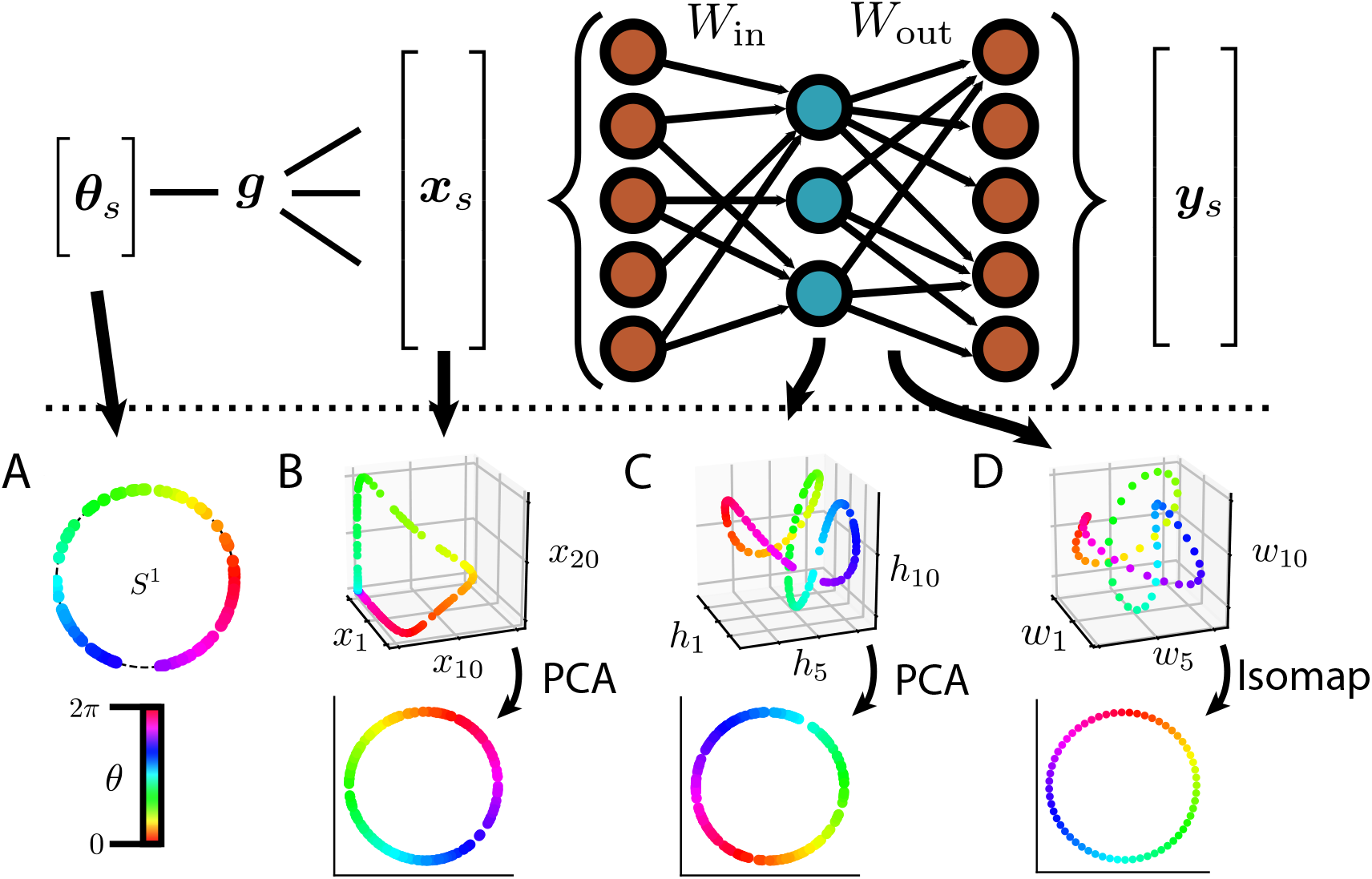
Overview of the characteristics of an autoencoder network trained to reproduce inputs that are generated by a periodic latent variable. Top: Network architecture. A low-dimensional set of latent variables is transformed into a high-dimensional input via a function ***g***. The network is then trained to reconstruct this input under the constraint of a bottleneck in the number of hidden unit neurons. A hyperbolic tangent nonlinearity is used for the hidden units. Bottom: Example network measurements for a periodic, scalar latent variable *θ*_*s*_. Network is trained with λ = 4*e*^−6^ and *m* = 60. (A-C) Color denotes value of *θ_s_*. (A) Latent variable *θ*_*s*_ drawn from *S*^1^. Color denotes value of *θ*_*s*_. (B) Responses of receptive field neurons 1, 10, and 20 to *θ_s_*. The receptive fields are periodic. Bottom: The response ensemble projected down to a two-dimensional space with PCA. (C) Responses of hidden units 1, 5, and 10 to *θ*_*s*_, out of 10 hidden units. Bottom: The hidden unit response projected down to a two-dimensional space with PCA. (D) Columns 1, 5, and 10 of the output weight matrix, out of 10 columns. Color corresponds to the receptive field neuron index. While the structure is similar to (C), the relative scaling of the axes is different, as evidenced by the grid lines. Bottom: The output weight matrix projected onto a two-dimensional space with Isomap.

To build some intuition, consider the characteristics of the model for *ϕ* equal to the identity, λ = 0, and zero bias terms. The loss function Eq. (2) is then the same as that of principal component analysis, except that here we do not require orthonormality of the input (output) weight rows (columns). Without this constraint, the model is highly *non-identifiable*: it is impossible to infer the value of the parameters (here ***W***_in_ and ***W***_out_) by sampling from the model. This is because

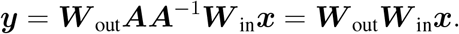

for any *N* × *N* invertible matrix ***A***, so that the same input-output mapping is satisfied by ***W***_out_***A*** and ***A***^−1^***W***_in_ for any invertible ***A***. When the orthonormality condition is enforced, the same statement holds excepting that ***A*** in this case is an arbitrary orthogonal matrix as opposed to an arbitrary invertible matrix. Since we are focused on investigating the structure of the weights, it is important to push the network toward finding a set of solutions that is unique up to orthogonal transformations, as these are geometry preserving. This is accomplished by the reqularizer term 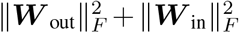 in Eq. (2) as we will discuss in more detail in Sec. 4.

Note that the matrix ***A*** that is found when training a neural network on the unregularized loss (λ = 0) may in practice not disturb geometric information very much. Specifically, if the Frobenius norm of ***A*** is not far from 1, then the geometric structure of ***W*** _out_***A*** compared to ***W*** _out_ will be similar, and similarly for ***W*** _in_. Choice of initialization of the weights before training, adding noise to the inputs while training, or implicit regularization caused by SGD may bias solutions toward regimes where the regularizer term of Eq. (2) is small. We have found experimentally that random weight initializations are often sufficient to lead to learned weights with recoverable structure even without regularization (data not shown).

We describe the training dataset next.

## 3 Constructing inputs generated by latent variables

### 3.1 Receptive field encoding of latent variables

Suppose that our inputs have the form ***x***_*s*_ = ***g***(***θ***_*s*_) ∈ ℝ^*m*^, where 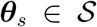 is some low-dimensional underlying process in a space 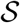 and ***g*** maps this process into a higher-dimensional space embedded in ℝ^*m*^. Here ***θ***_*s*_ is a latent variable that underlies the inputs ***x***_*s*_, and the induced process ***x***_*s*_ can be thought of as a high-dimensional encoding of ***θ***_*s*_. The subscript *s* is the index for samples of the corresponding variables, and is sometimes suppressed when context makes it clear. Throughout, mathematical symbols in bold font denote vectors or vector-valued functions, while scalars are denoted by lowercase symbols with normal font. When indexing the entries of these vectors with an index *k*, we either use the notation (*x*_*k*_)_*s*_ or suppress the *s*, simply writing *x*_*k*_. In our mathematical analysis and simulations we assume that the samples ***θ***_*s*_ are drawn uniformly from 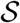.

The high-dimensional nature of the inputs is important in our framework. While networks that take lower-dimensional inputs and project them into a higher-dimensional hidden representation space are also of general interest, our objective here is to require the network to extract latent variables from a high-dimensional signal. This more constrained scenario affords a greater possibility that the weights will contain information about the latent variables. In particular, when treating weight matrices as geometric objects (as in Fig. 1D), the number of input dimensions *m* is the number of datapoints that we plot, and these points are embedded in an *N*-dimensional space. From this perspective, *m* will need to be large enough for meaningful structure to emerge.

One important class of encodings that increase the dimensionality of the encoded variables are the tuned neural response functions. In this case, each component *x*_*k*_ of ***x*** is the response of a neuron with tuning curve *g*_*k*_ to the variable ***θ***. As an example, consider the classic case of direction-selective retinal ganglion cells. We suppose each ganglion cell to be tuned as a Gaussian centered at its preferred orientation, *g*_*k*_(*θ*) ∝ exp(−*d*(*θ*, *z*_*k*_)^2^/*σ*^2^), where *θ* is an angle in the interval from 0 to 2*π*, *z*_*k*_ is the neuron’s preferred orientation, and *σ* captures the width of the tuning. Here *d* is a distance function in angle space, which can be written *d*(*θ*, *θ′*) = min{|*θ*−*θ′*|, 2*π* − |*θ*−*θ′*|}. A visualization of *θ* and the response ***g***(*θ*) is shown in Fig. 2A. The characteristics of an autoencoder network trained to reproduce this periodic input is shown in Fig. 1. Here we see that the periodicity of the inputs appears in both the hidden representation of the network as well as the weights. This will be elucidated in Sec. 4.

**Figure 2:**
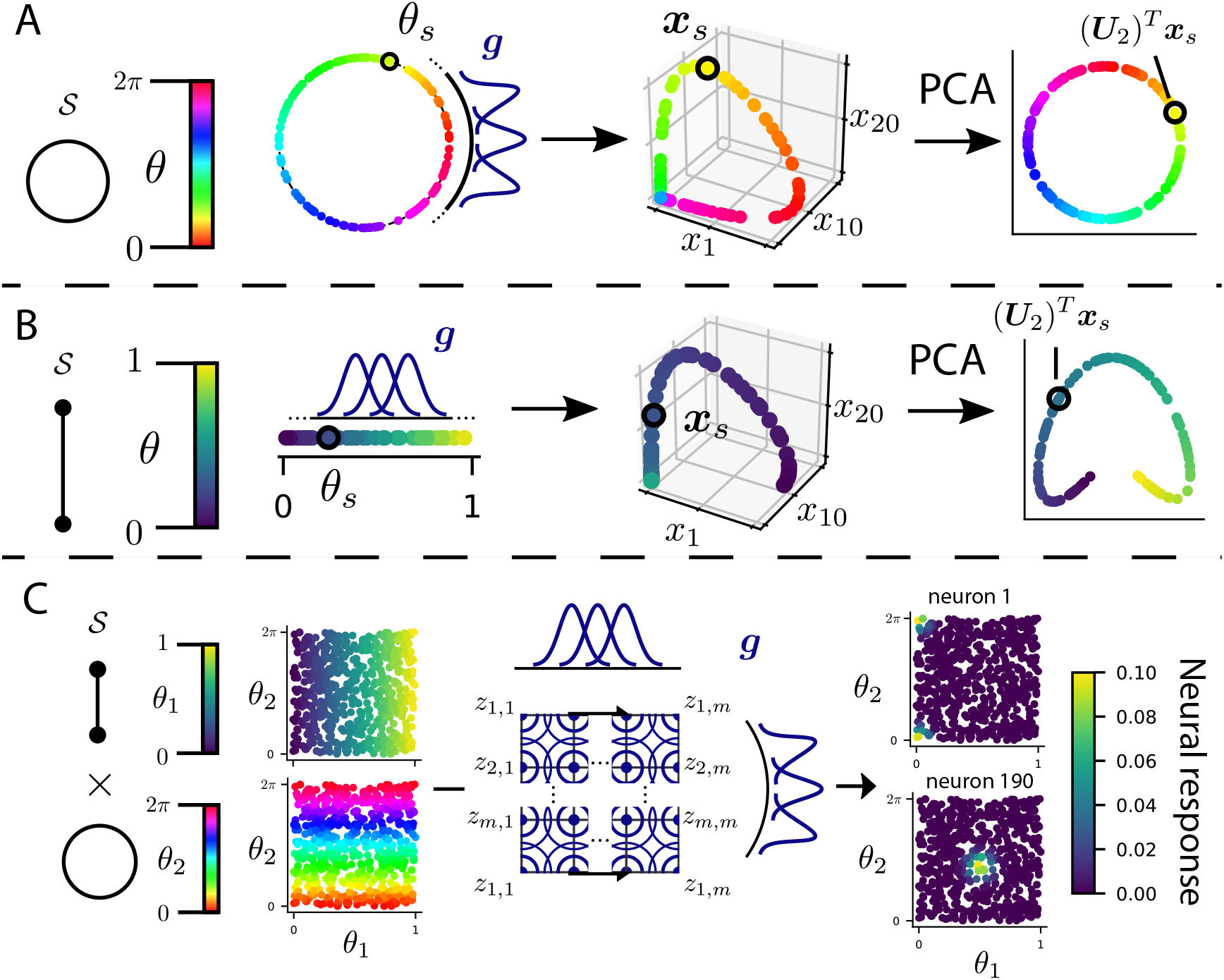
Depiction of latent variables on different spaces 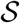. (A) Example where 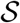 is a circle. A periodic scalar latent variable is transformed into a higher-dimensional encoding via the receptive field neural response function ***g***. The periodicity of *θ* is expressed by a periodic colormap for *θ*. The periodic structure is revealed by PCA. (B) Same as (A), but for a nonperiodic scalar latent variable, so that 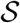 is a line segment.(C) Example where 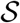 is a cylinder. Tensor product of a nonperiodic (*θ*_1_) and a periodic (*θ*_2_) latent variable is transformed into a higher-dimensional encoding via the receptive field neural response function ***g***. The scatter plots to the left depict the samples ***θ***_*s*_ with coloration based on *θ*_1_ (top) and coloration based on *θ*_2_ (bottom). Receptive field centers ***z***_*k*,*k′*_ tile the latent space 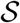. Specifically, the top and bottom edges of the space are glued together. The responses of the first and 190th out of 400 receptive field neurons are shown on the right.

Fig. 2B depicts a latent variable *θ* that is also scalar-valued, but differs from the previous example in that the latent space 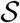 is the closed interval [0, 1] with the standard topology of an interval, as opposed to a circle. In this case the natural encoding is *g*_*k*_(*θ*) ∝ exp(−*d*(*θ*, *z*_*k*_)^2^/*σ*^2^) where *d*(*θ*, *z*_*k*_) = |*θ*−*z*_*k*_|.

Another important case is that of “place cell” neurons tuned to (*x*, *y*) position on a grid over a two-dimensional flat surface 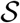. The space 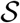 can be thought of as the product space of the product of two line segments. In this case we suppose each place cell to be tuned as a gaussian centered at its location on the grid, *g*_*k*,*ℓ*_(***θ***) ∝ exp(−*d*(***θ***, ***z***_*k*,*ℓ*_)^2^/*σ*^2^) where ***z***_*k*,*ℓ*_ is the neuron’s tuning center and *d*(***θ***, ***z***_*k*,*ℓ*_) = ‖***θ*** − *z*_*k*,*ℓ*_‖_2_. Note that in this case the indices (*k*, *ℓ*) need to be “unrolled” into a vector to form the vector-valued ***g***(***θ***_*s*_). The idea can be further extended to other latent variables, such as the joint spatial and orientation tuning seen also in retinal ganglion cells. To illustrate such a joint encoding, consider “place cell” neurons as before, but instead of tiling a grid, suppose that one edge of the grid wraps around and connects to the opposite edge, so that the neurons tile a cylinder. A realization of a latent variable on a cylinder and the responses of two receptive field neurons is depicted in Fig. 2C.

### 3.1 The latent variables appear in the weights of the trained autoencoder

Each of these examples illustrates a different topology of the space 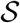 on which the latent variable ***θ***_*s*_ can live. Our main finding is that, in our network trained to autoencode the inputs ***x***_*s*_ generated by ***θ***_*s*_, the weights of the network generally reflect the topology of 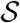. This is in addition to the hidden unit activations in response to the inputs reflect-ing the topology of 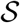. An overview of this phenomenon in the case of a periodic latent variable is given in Fig. 1 as well as in Figs. 3A to 3C. In Fig. 3A, a trained autoen-coder’s response to the inputs ***x***_*s*_ generated by a periodic latent variable *θ*_*s*_ is plotted in the top two-dimensional principal component space. In Fig. 3B, the top two principal components of the columns of the output weights are plotted. The resulting structure suggests periodicity, but isn’t always clearly seen. Using the nonlinear dimensionality reduction method Isomap [15] to reduce the output weights to a two-dimensional space reveals the circular structure of the latent variable space clearly (Fig. 3C). A description of Isomap and explanation of its success in recovering the periodic structure is given in Sec. 4.3. This shows that the structure of the latent variable of the trained autoencoder is apparent not only in the hidden unit activities, but also in the learned weights of the network.

**Figure 3:**
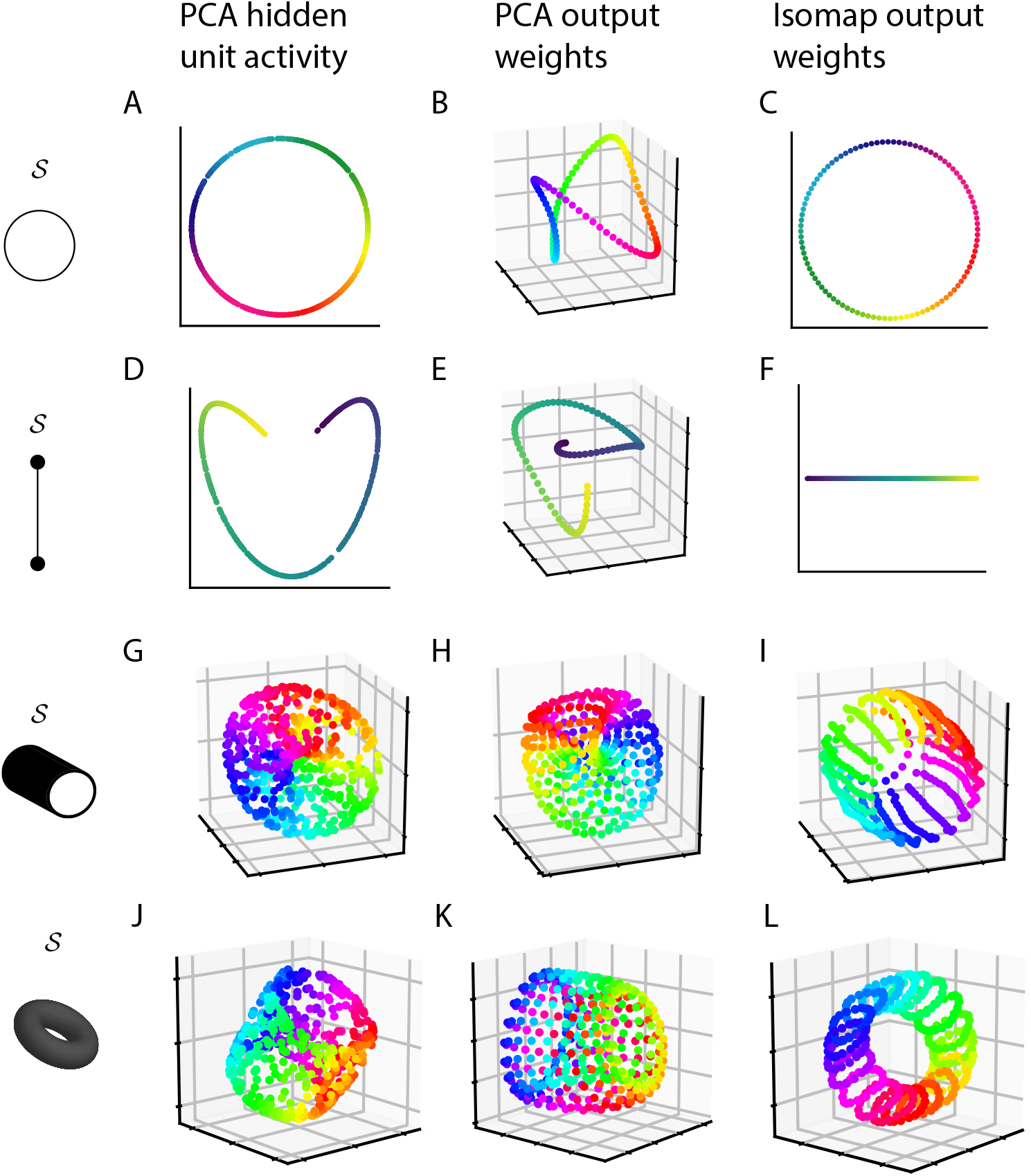
The structure of the latent variables can be recovered from the weights of the trained autoencoder by nonlinear dimensionality reduction methods. (A) Hidden unit activations of the autoencoder trained to reconstruct an encoding ***g***(*θ*_*s*_) of a periodic latent variable *θ*_*s*_ as in Fig. 2A and using the same coloration. (B) Principal components of the columns of the output weights of the autoencoder trained on the periodic latent variable. (C) Two-dimensional embedding via Isomap of the columns of the output weights for the network trained on the periodic inputs. (D-F) As in (A-C) but for an encoding ***g***(*θ*_*s*_) of a nonperiodic latent variable *θ*_*s*_ as in Fig. 2B. (G-I) As in (A-C) but for a joint encoding ***g***(***θ***_*s*_) of a periodic and non-periodic latent variable, such as that illustrated by Fig. 2C. In this case the Isomap embedding in (I) is three-dimensional. In (G) color corresponds with the periodic latent variable, while in (H-I) coloration is by the index of the receptive field centers corresponding to the periodic latent variable. (J-L) Same as in (G-I), but for a joint encoding of two periodic latent variables. Color corresponds with the first latent variable. The latent space 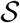 in this case is a torus.

A similar phenomenon occurs for an autoencoder trained to reconstruct inputs ***x***_*s*_ formed by receptive field responses ***g***(*θ*_*s*_) to a non-periodic latent variable *θ*_*s*_. Here we see that the topology of the latent space 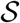 is again reflected in the network weights (Figs. 3E and 3F).

This phenomenon also occurs in the case of inputs generated by tensored latent variables as in Fig. 2C, resulting in the weights reflecting the cylindrical topology of 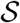 (Figs. 3G to 3I). When both tensored variables are periodic, the structure in the weights is that of a torus (Fig. 3L)

While in the examples above the latent variables are reflected both in the structure of the weights and the structure of the hidden layer activations of the trained network, structure in the activations depends on the choice of inputs given to the network after training. In the example of a periodic random variable, the ring structure in the activations does not appear if white noise inputs are shown to the trained network (data not shown). This illustrates how the information provided by the weights is in some ways distinct from that provided by the hidden unit activations.

Note also that the structure of the weights is most clearly extracted by the nonlinear dimensionality reduction method Isomap (Figs. 3C, 3F and 3I) as opposed to the linear method of principal component analysis (Figs. 3B, 3E and 3H). We shed light on why this is in Sec. 4.3.

## 4 Extracting latent variables from network weights

We now explain these observations through mathematical analysis. For ease of analysis, we consider a linear autoencoder model where *ϕ* is taken to be the identity. Our analysis involves three steps. The first is to relate the minimizers of Eq. (2) to the familiar solutions found by principal component analysis (PCA). The second is to resolve the problem of *non-identifiability* of the model, as introduced in Sec. 2. Once the minimizers of Eq. (2) have been related to the PCA solutions and the degeneracy of the solution space has been resolved, the third step in our analysis is to look more closely at the PCA solutions and to show that these solutions in fact encode the latent variable information.

### 4.1 Relating autoencoders to PCA

We start by rewriting the loss Eq. (2) in matrix form with λ = 0 and with *ϕ* taken to be the identity:

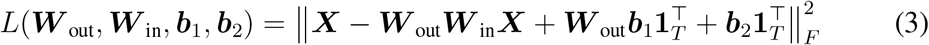

where ***X*** is an *m* × *T* matrix with column *s* holding the sample ***x***_*s*_ and **1**_*T*_ is the length *T* vector of all ones. Let ***μ***_*x*_ = 〈***x***_*s*_〉_*s*_ and ***μ***_*h*_ = 〈***h***_*s*_〉_*s*_. The optimal values for the bias terms have the effect of transforming the problem into one that has been mean-centered, i.e. the minimal weights of Eq. (3) coincide with those of

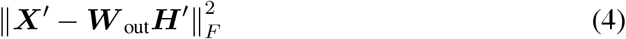

where ***X**′* = ***X*** − ***μ***_*x*_**1**^⊤^ and ***H′*** = ***W***_in_***X*** − ***μ***_*h*_**1**^⊤^ [16]. Minimizing this loss while enforcing that ***W***_in_ and ***W***_out_ have orthonormal rows and columns, respectively, results in the PCA solution. This solution is naturally expressed in terms of the *singular value decomposition* (SVD) of ***X**′*: ***X**′* = ***UΣV***^*T*^ where ***U*** is an *m* × *m* orthogonal matrix, ***V*** is a *T* × *m* matrix with orthonormal columns, and ***Σ*** is an *m* × *m* diagonal matrix with nonnegative entries *σ*_1_, *σ*_2_, …, *σ*_*m*_ called the *singular values* of ***X**′*. The standard PCA solutions are then 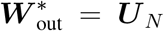 and 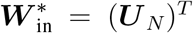 where ***U***_*N*_ is the matrix ***U*** truncated to the first *N* columns. However, as discussed above any solution of the form 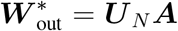 and 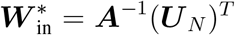 is also a global minimum, where ***A*** is an arbitrary invertible *N* × *N* matrix. This is the most general form of optimal solution [17, 18].

### 4.2 Resolving non-uniqueness of the optimal weights

The arbitrary invertible linear transformation ***A*** described in the previous section can potentially skew the structure of the weights past the point where the latent variables can be extracted. While ***A*** preserves topological information, it doesn’t necessarily preserve local distances between points. Nonlinear dimensionality reduction methods like Isomap are generally designed to embed points in a lower-dimensional space while preserving local distances, and losing information about these local distances can be destructive. This can become a problem in practice since the number of columns (rows) of ***W***_in_ (***W***_out_) are finite, and the structure of local distances among finitely many points can be lost as skewing becomes large. Here we describe biologically motivated conditions that address this issue.

If we add the regularizer 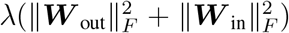 with λ > 0 to Eq. (3), then the solution becomes more constrained. Let **Σ**_λ_ be the diagonal matrix 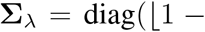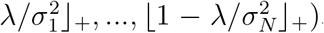, where the *σ*_*k*_ are again the singular values of ***X***, λ is the scaling of the regularizer, and ⎿·⏌_+_ is the threshold function max{·, 0}. [17, 18] recently showed that, under the assumption that *σ*_1_ > *σ*_2_ > … > *σ*_*N*_, the optimal weights in this case have the form 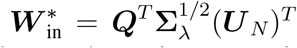 and 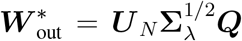 where ***Q*** is an arbitrary *N* × *N* orthogonal matrix. In particular, these solutions are unique up to arbitrary orthogonal transformations ***Q***, rather than arbitrary invertible transformations ***A***.

Therefore, this regularizer can help to preserve the geometric information encoded in the weights found by optimization methods. In particular, for positive but small λ, 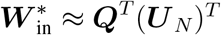 and 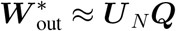. Since ***Q*** preserves distances (in other words, ***Q*** preserves geometric information), analyzing the geometric structure of optimal weights 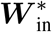 and 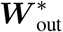 reduces to analyzing the geometric structure of ***U***_*N*_.

As we show next, the matrix ***U***_*N*_ contains geometric information about the inputs.

### 4.3 Relating the weights to the latent variables

In the previous section we showed how the optimal weights share the same geometric structure as 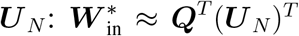 and 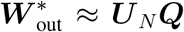 for λ small. We now show how ***U***_*N*_ is related to the latent variables underlying the inputs. To do so, we first note that according to basic properties of the SVD, ***U*** is a matrix of normalized eigenvectors of the covariance matrix 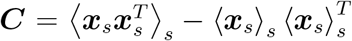 of ***X***, so **C** = ***U*****Σ**^2^***U***^*T*^ (recall that **Σ** holds the singular values for the mean-centered ***X′***). Assuming that ***x*** has the form *x*_*j*_ = *g*_*j*_(***θ***) ∝ exp(−*d*(***θ***, ***z***_*j*_)^2^/*σ*^2^), we can work out the form of the covariance between *x*_*j*_ and *x*_*k*_. Taking the limit *T* → ∞ and invoking the law of large numbers, we have that 〈(*x*_*j*_)_*s*_〉_*s*=1,…,∞_ = 〈*g*_*j*_(***θ***)〉_***θ***_ and 〈(*x*_*j*_)_*s*_(*x*_*k*_)_*s*_〉_*s*=1,…,∞_ = 〈*g*_*j*_(***θ***)*g*_*k*_(***θ***)〉_***θ***_. Letting *μ*_*j*_ = 〈*g*_*j*_(***θ***)〉_***θ***_, it follows that in this limit

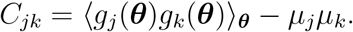

From this form of the covariance matrix, we can now work out the eigenvector structure for different choices of the latent space 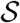 and distance function *d*.

#### Periodic latent variables give rise to periodic weight structure

We first consider the case of ap eriodic latent variable *θ*. Recall that the inputs are formed by encoding *θ* via orientation selective receptive fields *g*_*k*_(*θ*) ∝ (exp(−*d*(*θ*, *z*_*k*_)^2^/*σ*^2^), with *d* being a distance function in angle space, *d*(*θ*, *θ′*) = min{|*θ*−*θ′*|, 2*π* − |*θ* − *θ′*|}. Assume that the receptive field centers *z*_*k*_ evenly tile the space.

The salient structure of this encoding can be expressed through the idea of *equivariance*. Let 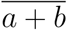 denote addition of *a* and *b* modulo the number, *m*, of receptive field neurons. In our scenario, equivariance means that shifting the identity of the receptive field neuron is the same as shifting the input to the receptive field neuron: 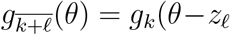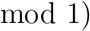. This equivariance implies a special structure of the input covariance matrix ***C***: ***C*** is a *circulant* matrix. This means that every row in ***C*** is a shifted version of the first row, where the shifting operation wraps around at the edges of the matrix. To show this, we show that entry *C*_*jk*_ is equal to 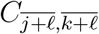, where *ℓ* is any integer.

Recall our assumption that *θ* is uniformly distributed on the circle 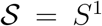, and suppose without loss of generality that 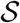 has Lebesque measure 2*π* (so *θ* varies from 0 to 2*π*). Then the probability density function of *θ* is the constant function that returns 1/(2*π*) for all *θ*. This means in particular that the expected value of *f* (*θ*) is 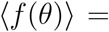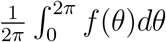 for any reasonably well-behaved function *f*.

To show that ***C*** is circulant, we first compute that

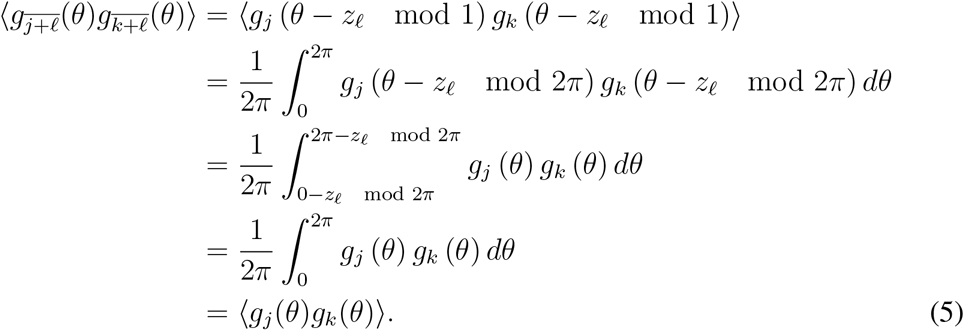

Computing shifts of the mean *μ*_*j*_ has a similar flavor:

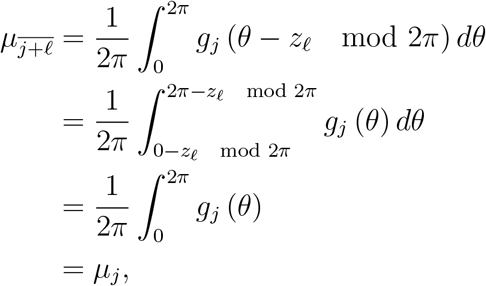

so that *μ*_*j*_ is independent of its index. Hence we can write *μ*_*j*_*μ*_*k*_ = *μ*^2^. Combining this with Eq. (5), we have that 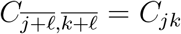. An example of the resulting circulant matrix is shown in Fig. 4A.

**Figure 4:**
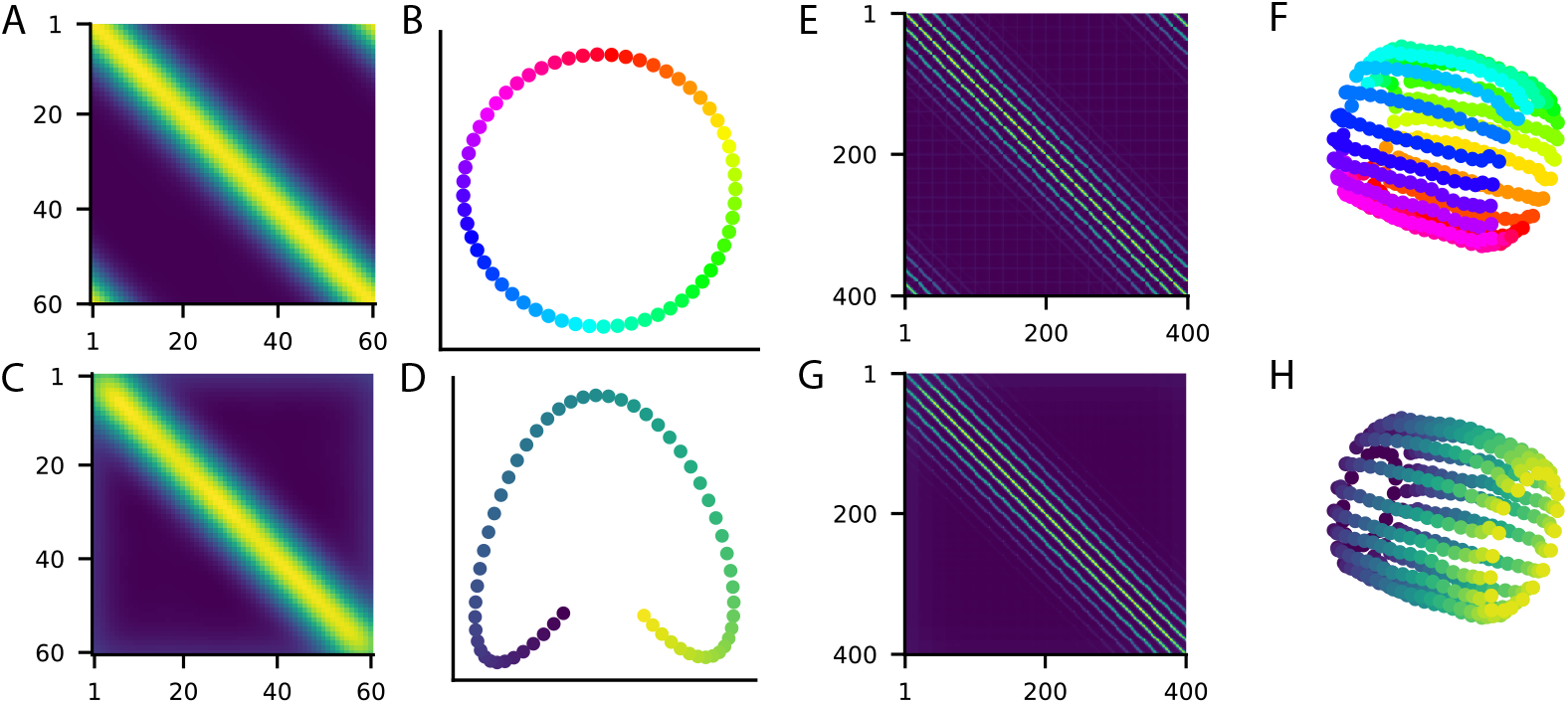
The structure of the latent variables can be recovered from the eigenvectors of the covariance matrix of the inputs. (A) Covariance matrix for the responses of *m* = 60 receptive field neurons to a periodic latent variable as in Fig. 2A. (B) Lowest frequency eigenvectors of the circulant matrix in A, plotted against each other and colored by index. (C) Covariance matrix for the responses of *m* = 60 receptive field neurons to a nonperiodic latent variable as in Fig. 2B. (D) Lowest frequency eigenvectors of the covariance matrix in (C), plotted against each other and colored by index. (E) Covariance matrix resulting from the tensored responses to a periodic and nonperiodic latent variable as in Fig. 2C, where the periodic variable is in the first coordinate and the nonperiodic variable is in the second. (F) Eigenvectors of the covariance matrix in (E), reduced to three dimensions by Isomap. Coloration is by the index of the receptive field centers corresponding to the periodic latent variable. The eigenvectors for the covariance matrix in (G) look similar. (G) Covariance matrix as in (E) but with the position of the nonperiodic and periodic variable switched. (H) Same as (F), but colored by the index of the receptive field centers corresponding to the nonperiodic latent variable.

In addition to being circulant, the covariance matrix ***C*** is by definition symmetric: ***C*** = ***C***^*T*^. The eigenvectors of circulant matrices are known and together make up the discrete Fourier transform matrix [19]. In the case where the circulant matrix is also symmetric, the real and imaginary parts of the eigenvectors are themselves eigen-vectors. This means that an eigenvector basis for ***C*** can be taken to be real, which results in eigenvectors that have one of three forms: cosine transforms, sine transforms, and the all-ones vector **1**_*m*_ (recall that *m* is the dimension of the inputs ***x***_*s*_). More precisely, the *j*th cosine transform eigenvector has the form 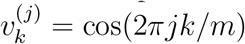 for *k* ∈ {0, 1, …, *m* − 1} and the *j*th sine transform eigenvector has the form 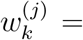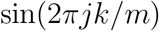. In particular, the eigenvectors are periodic, reflecting the periodicity of the latent variable *θ*. These eigenvectors together form the columns of the matrix ***U***. As an illustration, the eigenvectors ***v***^(1)^ and ***w***^(1)^ are plotted against each other in Fig. 4B.

Consider the truncation ***U***_*N*_ of ***U*** to *N* columns. We’re interested in the properties of ***U***_*N*_ embedded as a geometric object, with each row constituting a single data point in *N*-dimensional space. The sine and cosine structure of the eigenvectors ensures that this structure is periodic. In particular, the rows of ***U***_*N*_ are *m* samples from a loop that nonlinearly curves through *N*-dimensional space.

Since this loop structure of ***U***_*N*_ is nonlinearly embedded, nonlinear dimensionality reduction methods are well suited for recovering this structure. Indeed, since the singular values of ***U***_*N*_ are all 1, trying to extract structure from it with the linear method of PCA will only return a random set of the columns of ***U***_*N*_. In general, the ability of nonlinear dimensionality reduction methods to successfully extract the structure of interest from a dataset depends on having enough datapoints, and our situation is no exception. In our case, the number of datapoints is *m*, and this will need to be a large enough number for the dimensionality reduction method to succeed. The precise number of datapoints needed will depend on the specifics of the dimensionality reduction method used. To proceed, we will assume that *m* is sufficiently large.

For intuition as to why nonlinear methods work, we focus on the approach of the nonlinear method Isomap. The first step of Isomap involves building a graph on the datapoints where points that are sufficiently c lose a re c onnected b y a n e dge. Let’s consider the strategy of connecting every point to its two nearest neighbors. Then in our case this graph will indeed be a loop through high-dimensional space, and embedding this graph in two dimensions in a way that best preserves distance information reveals a ring.

Recall the minimizer 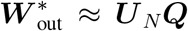, 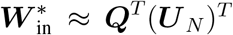 of the regularized loss for the linear model. In practice we find that the periodicity of ***U***_*N*_ is reflected in the weights ***W***_out_ in the autoencoder trained with stochastic gradient descent. As illustrated by Fig. 3, this extends to networks with tanh nonlinearity. Here the network is trained with λ = 4*e*^−6^ and *m* = 100. The latent variable structure can be partially seen in the apparent periodicity of points obtained by using PCA to project the columns of ***W***_out_ onto a three-dimensional space in Fig. 3B. As discussed above, this periodicity is revealed more clearly by using Isomap to “unravel” the coils caused by the higher frequency modes, as can be seen in Fig. 3C.

#### Nonperiodic latent variables give rise to nonperiodic weight structure

The above analysis can be repeated in a similar form for the case of a nonperiodic latent variable *θ* on a line segment, where this time *g*_*k*_(*θ*) ∝ exp(−|*θ*−*z*_*k*_|^2^/*σ*^2^). Suppose the receptive field centers *z*_1_ through *z*_*m*_ evenly tile the line segment [0, 1], with *z*_1_ = 0 and *z*_*m*_ = 1. While we are interested in the case where *θ* is uniformly distributed on [0, 1], this becomes mathematically challenging to work with due to conditions at the boundary being different than conditions in the center of the interval. Instead, we let 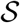 be the interval [−*s*, *s* + 1], with the usual interval topology. Taking *s* sufficiently large will allow us to deal with boundary effects; for instance, this assumption ensures that *g*_*k*_(*θ*) is approximately independent of *k*. In this case the covariance matrix, instead of being circulant, is approximately *Toeplitz*, which means that the entries on each descending diagonal from left to right are the same. This can be seen by choosing indices *j*, *k* and *ℓ* constrained such that *j*, *k* ∈ {1, …, *m*}, *j* + *ℓ* ∈ {1, …, *m*}, *k* + *ℓ* ∈ {1, …, *m*} and computing

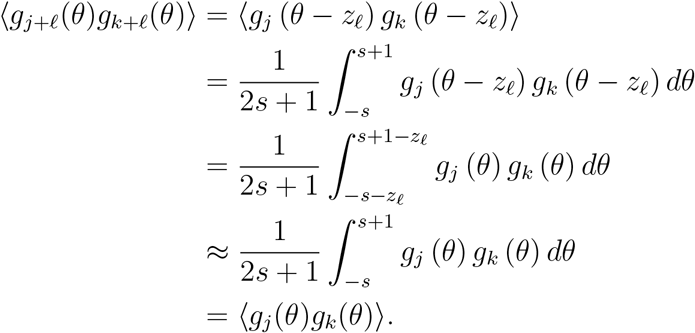

The approximation is justified when *s* is much larger than *z*_*m*_, since the contribution to the integral near the integration limits is vanishingly small. This approximation becomes exact for large enough *s* if we clip the receptive field functions *g*_*j*_ to have finite support.

Computing the shifted means has a similar flavor:

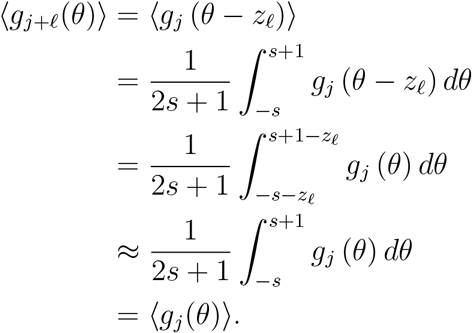

Taken together, these equations imply that *C*_*j*+*ℓ*,*k*+*ℓ*_ = *C*_*j*,*k*_, so that ***C*** is Toeplitz. In our simulations we take *s* = 0 so that ***C*** is only approximately Toeplitz, but find that the conclusions below still hold in practice.

While the eigenvectors of Toeplitz matrices are not in general determined as they are for circulant matrices, they are known for tridiagonal Toeplitz matrices. Symmetric tridiag nal Toeplitz matrices all have the same eigenvectors, of the form 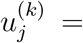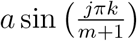 for *k*, *j* = 1, …, *m*, where *a* is an arbitrary nonzero scalar. The odd eigen-vectors ***u***^(2*k*+1)^ are symmetric (which in particular means that 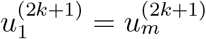) while the even eigenvectors are antisymmetric (in particular, 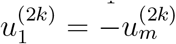). Recall that the ***u***^(*k*)^ make up the columns of ***U***.

As before, we consider ***U***_*N*_ as a geometric object embedded in *N* dimensional space, where the rows are datapoints. Under the assumptions of tridiagonal covariance matrix, the eigenvectors ***u***^(*k*)^ given above reveal a particular structure: in our numerical tests, the rows of ***U***_*N*_ lie along a curve with the endpoints disconnected, provided that *N* < *m*. To show this, we need to show that the distance the first and last row of ***U***_*N*_ is larger than the distance between adjacent rows. While we do not prove this here, in practice we have found this to be the case numerically (data not shown). In fact, for *m* − *N* sufficiently large we find that the distance between the first and last row is larger than the distance between rows *k* and *k* + 2, for *k* = 1, …, *m* − 2 as well.

With this “gap” between the first and last row of ***U***_*N*_, we can use nonlinear dimensionality reduction methods to reveal the structure of a line segment. Consider again the strategy of connecting every point to its two nearest neighbors, as is done in using Isomap. In this case the “middle” sections of the curve will look as in the case of the periodic latent variable, but the ends will be different. If *m* − *N* is large enough then the two endpoints of the line will not be connected, and the general structure of the graph will be that of a line.

Now we consider Toeplitz matrices with more than three (but still finitely many) nonzero diagonals. For fixed *k*, the eigenvector ***u***^(*k*)^ in this case approaches the form 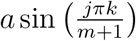 in the asymptotic limit of large *m* [20]. It follows that, after truncating to ***U***_*N*_ for finite *N* and taking *m* to be large, we can use the same reasoning as in the tridiagonal case to infer that the rows of ***U***_*N*_ lie along a curve with the endpoints disconnected. The structure of the eigenvectors is illustrated by plotting ***u***^(1)^ and ***u***^(2)^ against each other in Fig. 4D for *m* = 60.

This topology appears in the weights of the trained autoencoder, as shown by Isomap in Fig. 3F. Here the network is trained with λ = 4*e*^−6^ and *m* = 100. PCA projections of the weights do not reveal this structure as clearly (Fig. 3E).

#### Tensored latent variables give rise to tensored weights

In this section we consider combinations of latent variables found by taking tensor products of other latent variables. Consider the case of “place cell” encoding on a torus, where both boundaries of the grid are periodic. This can be thought of as a tensored combination of two periodic latent variables. Suppose that the first and second coordinates of ***θ*** correspond to the periodic latent variables *θ*_1_ and *θ*_2_, respectively, and that each is i.i.d. uniformly distributed on the circle *S*^1^.

Recall our choice of Gaussian curve response function on the circle: 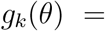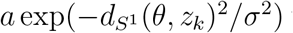 where *a* is a positive scalar and 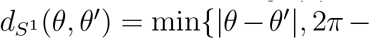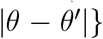 is distance on the circle. We abuse notation slightly and use the same name for a Gaussian curve response function on the torus: *g*_*i*,*k*_(***θ***) = *a*^2^ exp(−*d*(***θ***, ***z***_*i*,*k*_)^2^/*σ*^2^) where *d* is Euclidean distance on the torus, which can be written

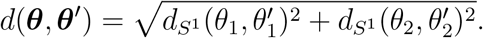

This time the tuning curve centers ***z***_*i*,*k*_ have two indices and evenly tile the two-dimensional surface of the torus. Our goal is to decompose ***C***_*i*,*j*,*k*,*ℓ*_ = 〈*g*_*i*,*k*_(***θ***)*g*_*j*,*ℓ*_(***θ***)〉−〈*g*_*i*,*k*_(***θ***)〉〈*g*_*j*,*ℓ*_(***θ***)〉 into contributions from tuning curves *g*_*k*_(*θ*) defined on the circle. We start with the observation that

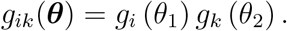

Using this, along with independence of *θ*_1_ and *θ*_2_,

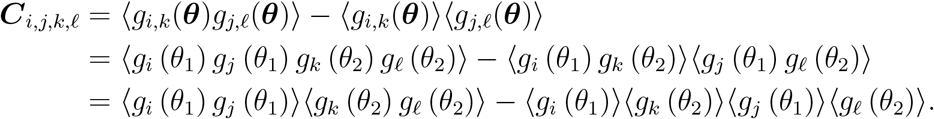

Recall that 〈*g*_*j*_(*θ*)〉 = *μ* is independent of *j*. If we let 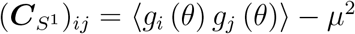 be the covariance matrix for inputs on a circle, then we can write

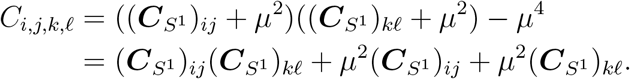

From this equation, we can see that ***C*** can be written as sums of Kronecker tensor products (denoted ⊗):

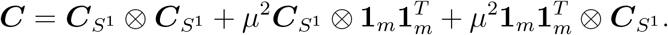

Matrix multiplication of Kronecker products satisfies the *mixed-product property*: (***A*** ⊗ ***B***)(***C*** ⊗ ***D***) = (***AC***) ⊗ (***BD***). Suppose that ***u*** and ***v*** are eigenvectors of 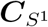 with eigenvalues λ_*u*_ and λ_*v*_, respectively. Then

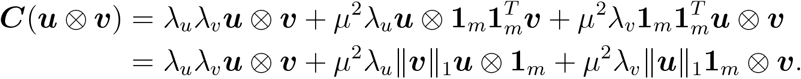

This equation reveals that ***u*** ⊗ ***v*** are eigenvectors of ***C*** provided either (1) *μ* = 0 or (2) ‖***u***‖_1_ = 0 or ***u*** = **1**_*m*_, and ‖***v***‖_1_ = 0 or ***v*** = **1**_*m*_. The eigenvectors of 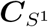 satisfy condition (2), as is easy to verify. Hence in the case of inputs formed from tensoring two periodic latent variables, we can find closed form solutions for the eigenvectors of the covariance matrix. These eigenvectors are tensor products of periodic latent variables, so that their structure reflects that of a torus (twisted nonlinearly through *N* dimensional space). This structure appears in the weights of the trained autoencoder (Fig. 3L). Here the network is trained with λ = 4*e*^−6^ and *m* = 20 · 20 = 400.

In the case where one or both of the variables being tensored is nonperiodic, we currently lack a general mathematical characterization of the eigenvectors. In the special case when the mean response is zero, *μ* = 0, the eigenvectors of ***C*** are the tensor products of the eigenvectors of the covariance matrices for the two latent variables. This can be shown by similar reasoning as above. Even when *μ* is nonzero, we see experimentally that the structure of the tensor product of a periodic and nonperiodic latent variable resembles a breaking of the toroidal structure similar to the scalar case. In particular, one end of the torus has a gap, which makes the structure resemble that of a cylinder. In this case the covariance matrix has the form shown in Fig. 4E if the periodic variable is the first coordinate and Fig. 4G if the nonperiodic variable is the first coordinate (this relationship may be reversed depending on how the four indices of ***C*** are unrolled into two indices). The cylindrical structure can be seen in Figs. 4F and 4H. This cylindrical structure is also reflected in the weights of the autoencoder (Fig. 3I). Here the network is trained with λ = 4*e*^−6^ and *m* = 20 · 20 = 400.

## 5 Discussion

It is important to investigate the ways that connectivity data can be used to help us understand neural circuits. Here we focus on using dimensionality reduction techniques to infer elements of the function of a neural circuit from the structure of the weights. We find that the latent variables that underlie the inputs can be recovered from the weights of an autoencoder with a single layer of hidden units. This is accomplished via nonlinear dimensionality reduction methods, such as Isomap. In particular, periodic inputs give rise to periodic weight structure, and nonperiodic inputs to nonperiodic weight structure. The tensor products of such inputs results in an analogous structure in the weights. The emergence of this structure depends on regularization to penalize large weights. It also depends on the inputs encoding low-dimensional latent variables in a high-dimensional way.

The approach of focusing on connectivity data to deduce information about the function of a neural circuit complements other very fruitful efforts of probing the activity of neurons in the circuit. The latter includes the seminal work of Hubel and Wiesel [21], which provided strong evidence via recordings in cat striate cortex that neural responses in this area are built from simple combinations of the responses of retinal ganglion cell neurons. Another noteworthy example is the analysis of bump-attractor-like dynamics in the *Drosophila* ellipsoid body [22], which demonstrated through two-photon calcium imaging that the circuit tracks orientation information through integrated sensory information. There are, however, difficulties in using neural activations alone to draw inferences. For instance, it isn’t always clear how to satisfactorily explore the space of all possible input stimuli. Often multiple competing models arise to reproduce neural circuit function or neural activity, and connectivity data can be used to select among them [6]. Connectivity data may also be useful for choosing parameters in models that are overparametrized. In the *Drosophila* ellipsoid body example, fine-grain analysis of connectivity data will probably be necessary to answer once and for all whether the bump attractor dynamics are implemented by a ring attractor network topology, and how this ring attractor is implemented precisely (see [23] for significant recent steps in this direction).

As analysis of connectivity data has its own set of shortcomings – for instance, information about neural modulation, the precise nonlinear responses of neurons to inputs, and many other factors are left out – hybrid methods that take into account both neural activation as well as connectivity data will be important far into the future. There are also other promising avenues for using connectivity data in ways distinct from the methods considered here. One approach is to develop models that fit neural activity data or that satisfy the believed function of a neural circuit while constraining them with connectivity data (reviewed in [6]). Another fruitful approach is to first generate a network model with a connectivity determined through data-driven means, assume a form of neural unit dynamics in the model, and then analyze the resulting network dynamics (for instance, [24, 25]).

Our approach is limited both by the simplicity of the task and network model, as well as the need to have exact values for the weights of synapses between neurons (a value that is difficult to assign in data). Our mathematical analysis assumes linear neural responses, and while we observe that these results extend in this case to hyperbolic tangent nonlinear responses in simulations, more work needs to be done to see how robust these approaches are to the type of nonlinearity used.

Our analysis opens the door to many interesting future studies. These include extensions to more complicated tasks and models such as deeper autoencoders or more general feedforward networks trained on more sophisticated tasks. It would also be valuable to determine if the structure can be recovered when exact synapse values are not known, when sparsity constraints on the weights are applied, or when different nonlinearities are used in the model. Brain circuits contain both deep hierarchy and recurrent connections, and it remains to be seen if our methods will be successful in artificial networks that have these complexities. In extending to data from the brain, persistent homology techniques could potentially be combined with nonlinear dimensionality reduction techniques to help deal with inaccuracies in the data.

The observation that using nonlinear – as opposed to linear – dimensionality reduction methods is important for extracting structure in the weights, and that regularization during training also encourages this structure to emerge, can guide efforts to investigate more complex models. In general, network models fail to be identifiable, exemplified by the arbitrary invertible matrix *A* in our model. In the same way, in deep linear networks arbitrary invertible linear transformations of one layer’s weights can be undone by the inverse transformation applied to the next layer’s weights. When it comes to analyzing neural networks (be it the connectivity or unit activities), it is important to work out the most natural constraints that result in *meaningful* and *interpretable* network structures. Here we’ve shown that the solutions enforced by Frobenius norm regularization [17, 18] are sufficiently constrained to yield latent variable information. This regularization can be viewed as a cost on weight resources, a biologically relevant constraining factor. This indicates that biological connectivity data may indeed be constrained such that they yield information about neural circuit function via dimensionality reduction methods like those explored here. In addition, we’ve found that this regularization is not always necessary for extracting the weights when training with SGD (data not shown). This may be because, with the right initialization, the solutions found by SGD are biased to have relatively low Frobenius norm. It is still an open question as to if L2 regularization or other constraints are sufficient for enforcing interpretability in broader classes of network models (see [14, 26, 27] for works related to these issues).

## 6 Acknowledgements

MF is funded by the National Science Foundation Graduate Research Fellowship under Grant No. DGE-1256082. ESB acknowledges the support of NSF DMS Grant 1514743. We thank the Allen Institute for Brain Science founders, Paul and Jody Allen, for their vision, encouragement, and support.

